# Transcriptome analysis reveals *CCND2* promotes myogenic differentiation of pig skeletal muscle

**DOI:** 10.64898/2026.09.16.752030

**Authors:** Jinbao Li, Jianmin Zhang, Qi Song, Shiyin Li, Yongqing Zeng, Wei Chen

## Abstract

**Objective:** Skeletal muscle growth rate differs substantially among diverse pig breeds. In the present study, RNA-seq was performed to identify differentially expressed genes (DEGs) across pig breeds with distinct growth rates. *CCND2* showed obvious differential expression among pig breeds. Although *CCND2* is essential for cell cycle control, its biological function in skeletal muscle differentiation has rarely been reported. This work aims to identify DEGs in pig breeds with different growth rates and explore *CCND2*’ role in porcine skeletal muscle development.

**Methods:** RNA-seq was performed to identify differentially expressed genes (DEGs) across pig breeds with distinct growth rates. Weighted gene co-expression network analysis (WGCNA) was used to identify gene modules potentially associated with skeletal muscle growth and development. Quantitative reverse transcription polymerase chain reaction (qRT-PCR) analysis was performed to validate the role of *CCND2* in skeletal muscle differentiation.

**Results:** We identified 5,416, 4,846, and 2,777 DEGs in the Duroc pig (DU) vs. Jiangquan black pig (JQ), DU vs. Laiwu pigs (LW), and JQ vs. LW comparison groups, respectively. WGCNA identified two gene modules (lightgreen and darkmagenta) potentially associated with skeletal muscle growth and development. *CCND2* exhibited differential expression in comparisons between DU vs JQ and DU vs LW, with higher expression observed in Duroc pigs. qRT-PCR analysis revealed that overexpression of *CCND2* at day 6 (D6) of cell differentiation significantly increased the expression levels of *CCND2* gene and markers of myogenic differentiation (*p* < 0.05). Conversely, knockdown of *CCND2* significantly decreased the expression levels of *CCND2* gene and myogenic differentiation markers (*p* < 0.05).

**Conclusions:** This study identified breed-specific DEGs and key gene modules in porcine skeletal muscle and verified that *CCND2* promotes myogenic differentiation. These findings provide a genetic theoretical basis for further investigations into skeletal muscle growth and development.

## INTRODUCTION

Skeletal muscle is a heterogeneous tissue composed of muscle fibers and connective tissue, primarily responsible for body movement through contraction and relaxation [1]. Skeletal muscle constitutes the predominant tissue in pigs, accounting for 45%–60% of total body weight. Its developmental status profoundly affects meat yield and largely defines porcine growth performance [2]. Beyond its locomotor function, skeletal muscle also can act as a critical regulator of the body’s protein reserves, glucose metabolism, and lipid homeostasis [3,4]. Abnormal growth of skeletal muscle is linked to various pathological conditions, including Duchenne muscular dystrophy [5], spinal muscular atrophy [6] and cancer cachexia [7]. Improving growth rate and meat production performance is a primary goal in pig breeding. The core task of the swine industry is to achieve maximum pork production with minimal investment while simultaneously enhancing pork quality. Research on porcine skeletal muscle is crucial for advancing animal husbandry, improving food quality, and safeguarding human health.

Jiangquan black pig is a crossbred based on the Yimeng black pig with introgression of Duroc pig which accounts for about 35% of the overall genetics. This breed not only retains the main advantages of the Yimeng Black pig, including distinct phenotypic characteristics, high disease resistance, tolerance to roughage, and superior meat quality, but also inherits desirable traits such as rapid weight gain and high lean meat percentage, thus possessing significant economic value [8]. Duroc pigs are globally renowned for their fast growth rate, high feed conversion efficiency, and high lean meat percentage [9,10], making them a widely recognized lean-type pig breed worldwide. As a superior pig breed native to North China, Laiwu pigs feature abundant intramuscular fat deposition [11,12], yet they exhibit a relatively slow growth rate.

In recent years, with the rapid development of biological sciences and sequencing technologies, research on the transcriptome of pig skeletal muscle has also made significant progress. Previous studies have demonstrated that bioinformatics analysis of miRNA and mRNA transcriptomes in the skeletal muscles of Changbai pigs, Tongcheng pigs and Wuzhishan pigs has identified differential expression of miRNAs and mRNAs among these breeds; furthermore, these differentially expressed genes are enriched in biological processes (BP) related to skeletal muscle growth and development [13]. Another study performed sequencing analysis on skeletal muscles from wild-type Meishan pigs and Meishan pigs with ZFN-edited myostatin deletion, identifying 438 DE genes. The above observations lay a solid theoretical foundation for exploring the regulatory mechanism of porcine skeletal muscle development, and contribute to the high-efficiency screening and recognition of candidate genes controlling pig growth characteristics [14]. Additionally, a study conducted whole-transcriptome sequencing on Duroc pigs with different average daily gains, identifying key lncRNAs and mRNAs that regulate late-stage growth and development in large-sized pigs, which may be critically involved in growth traits and skeletal muscle development [15].

The CCND family genes, including *CCND1*, *CCND2*, and *CCND3*, were first isolated from thyroid cancer [16]. These three D-type cyclins exhibit high homology [17]. As the regulatory component of Cyclin D2-CDK4 heterodimer, *CCND2* triggers phosphorylation and functional repression of RB family proteins represented by RB1. Phosphorylated RB1 loses its binding capacity to E2F transcription factor, releasing E2F to activate the transcription of its target genes. These gene products are essential for cells to complete the G1 phase of the cell cycle [18]. At present, studies on *CCND2* have mainly focused on cancer and the reproductive process [19–22]. However, emerging evidence indicates that *CCND2* also participates in skeletal muscle growth and development. A previous study reported that MyoD1 regulates *CCND2* expression via the PI3K-Akt signaling pathway, thereby contributing to myogenesis and muscle differentiation in Guangling cattle. *MyoD1* knockout significantly downregulates *CCND2* expression, suggesting that *CCND2* functions as a crucial downstream regulator during muscle cell differentiation [23]. Nevertheless, direct evidence supporting the role of *CCND2* in skeletal muscle differentiation remains limited.

Accordingly, further investigation is warranted to clarify the function of *CCND2* during skeletal muscle differentiation.

The present study focused on three pig breeds with distinct growth rates. RNA-seq was performed on longissimus dorsi muscle samples from eight Jiangquan Black pigs, and transcriptome datasets for eight Duroc pigs and nine Laiwu pigs were downloaded from the NCBI database. Bioinformatics analyses were conducted to screen core genes associated with skeletal muscle growth and development, and *CCND2* was selected for further functional validation in C2C12 myoblast differentiation. This study aimed to identify DEGs among various pig breeds and elucidate the role of *CCND2*, thereby improving pig growth performance and providing a genetic basis for future studies on skeletal muscle growth and development.

## MATERIALS AND METHODS

### Ethics Statement

The animal study was reviewed and approved by the Animal Ethics Committee of Shandong Agricultural University, China, and performed in accordance with the Committee’s guidelines and regulations (SDAUA-2023–157).

### Sources of animal tissue samples and raw data

Jiangquan Black pigs come from Shandong Linyi Jiangquan Agricultural and Animal Husbandry Co., Ltd. Eight Jiangquan Black pigs were fed under identical management, conditions, and nutritional standards. Prior to slaughter, the pigs were electrically stunned to induce immediate unconsciousness and then euthanized by exsanguination via severance of the carotid artery. After slaughter, the longissimus dorsi muscle was swiftly obtained at the final thoracic vertebra of the carcass for subsequent total RNA extraction. The raw data of Duroc pigs and Laiwu pigs were obtained by downloading from the SRA database using sratoolkit (version 3.0.0). The SRA number for Duroc pigs was PRJNA812354, while the SRA numbers for Laiwu pigs were PRJNA815878 and PRJNA354523. All samples were collected from longissimus dorsi muscle of adult pigs and were suitable for comparative transcriptomic analysis.

### RNA Extraction, Library Construction and Sequencing

Longissimus dorsi muscle RNA was extracted with TRIzol (Invitrogen) and quality-checked on an Agilent 2100 Bioanalyzer. Library construction entailed oligo(dT)-based mRNA capture, chemical fragmentation, cDNAsynthesis (both strands), end-repair, A-tailing, adaptor ligation, AMPure XP bead-based size selection, and PCR enrichment. The resulting libraries were subjected to PE150 sequencing on an Illumina NovaSeq 6000 platform (Biomarker Technologies, Beijing).

### Quality Control for Raw Reads and Mapping

Raw sequencing reads underwent quality filtering prior to genome mapping. Fastp [24] (version 0.23.2) with standard settings was adopted to trim raw data, eliminating adapter-contaminated sequences, and low-quality fragments. The genome sequence file used in this study for pigs was Sus_scrofa.Sscrofa11.1.dna.toplevel.fa. HISAT2 [25] (version 2.2.1) was applied to map clean reads of every sample to the Sus scrofa reference genome (Sscrofa11.1.fa).

### Quantification of gene expression and differential expression analysis

Gene read counts of aligned reads was subsequently performed using FeatureCounts [26] (version 2.0.3) with the corresponding gene annotation file (Sus_scrofa.Sscrofa11.1.108.chr.gtf.). FPKM values were calculated to normalize gene expression levels for sequencing depth, and DESeq2’s size-factor normalization was applied to correct for library size variation. Differential expression analysis on biologically repeated samples was conducted with DESeq2 [27] (version 1.34.0). Benjamini–Hochberg correction was applied to adjust raw *p* values and limit false discovery rate. Genes satisfying FDR < 0.05 and |log2(Fold Change)| ≥ 1 were regarded as DEGs.

### Weighted Gene Co-expression Network Analysis

WGCNA [28] characterizes gene correlation patterns across distinct specimens. This method screens gene clusters that exhibit high levels of co-expression, analyze the correlation between modules and specific features or phenotypes. Currently, this method has been widely employed in studies associating phenotypic features with related genes, among other research areas. This analytical method differs from solely considering differentially expressed genes. In WGCNA analysis, thousands of genes are considered as input, aiming to identify modules of interest and correlate them with traits. Genes with absolute FPKM values over 0.3 were screened as analytical inputs in the current research, yielding a total of 7938 genes. The main steps of WGCNA are as follows: 1) Selecting the optimal sftpower; 2) Hierarchical clustering trees display various modules, correct and merge modules, and plot module hierarchical clustering diagrams; 3) Correlations between individual modules and traits were calculated, and gene modules of interest were selected and analyzed for GO enrichment and Kyoto Encyclopedia of Genes and Genomes (KEGG) pathway; 4) Visualize gene networks and display the correlations among modules.

### Gene ontology and Kyoto encyclopedia of genes and genomes enrichment analyses

GO and KEGG enrichment analyses were carried out on DEGs and key module genes to explore their biological roles. The GO database classifies gene functions into three hierarchies: biological process, cellular component (CC) and molecular function (MF). All DEGs and target module genes were submitted to DAVID 2021 [29]. GO terms and KEGG pathways with *p* < 0.05 were considered significant by the DAVID database. To visualize the results of the enrichment analysis, DAVID analytical outcomes were imported into SangerBox (3.0) and Hiplot Pro (2.0). The Circos plot represents the results of the DEGs analysis, while the Bubble plot represents the results of WGCNA module genes.

### qRT-PCR

GAPDH served as reference gene in qRT-PCR verification. Six DEGs were randomly picked to validate expression trends matching RNA-seq data. Primer Premier 5.0.0 was used to design corresponding primer pairs, which were synthesized by Accurate Biotechnology Co., Let, Changsha, China. For the functional experiment of the *CCND2* gene, β-actin was selected as internal control. Primers of *CCND2* and myogenic markers (*MyoG*, *MEF2C*, *Myf5*) were designed via the same software and synthesized by the same company. Amplification was performed on Roche Light Cycler 96. Relative gene expression was calculated with the 2^−ΔΔCT^ method [30]. The qRT-PCR primer sequences were showed in Supplementary Table S1.

### Overexpression and Knockdown Assay

For overexpression assay, full coding sequence of mouse *CCND2* (NM_009829.3) was generated, and high fidelity was ensured by using polymerase Gflex (TaKaRa, Dalian, China). Following purification, PCR amplicons were subjected to dual restriction digestion using Hind III and Xho I. The digested inserts were subsequently ligated into linearized pcDNA3.1(+) backbone with T4 DNA ligase. The resulting recombinant plasmids were chemically transformed into DH5α competent cells supplied by TaKaRa. All positive recombinants were confirmed via two-directional sequencing, followed by endotoxin-free midi plasmid purification prior to transfection into C2C12 myoblasts. The plasmid that was acquired was called pcDNA3.1(+)-CCND2. The control was an empty pcDNA3.1(+) vector.

The murine *CCND2* gene was used to guide the creation of siRNA for the knockdown experiment (Accurate Biotechnology Co., Let, Changsha, China). The siRNA negative control exhibited identical composition to the siRNA sequence; however, it lacked homology with the *CCND2* mRNA. Four siRNA were synthesized (Supplementary Table S2). The siRNA with optimal silencing efficiency was selected to investigate *CCND2* knockdown phenotypes.

### Cell Culture, Treatments and Transfection

C2C12 cells were maintained in culture flasks with complete DMEM medium containing 10% fetal bovine serum and 2% penicillin-streptomycin mixture. Cell incubation was performed at 37 °C under a humidified atmosphere with 5% CO. To trigger myogenic differentiation, fully confluent C2C12 cells at the contact inhibition state were cultured in differentiation medium consisting of DMEM supplemented with 2% horse serum, which was defined as Day 0 (D0). The differentiation medium was refreshed daily until Day 8 (D8). For transfection assays, passaged C2C12 cells were seeded in six-well plates. When cell density reached 70%–80% confluence, cells were transfected with pcDNA3.1(+)-CCND2 overexpression plasmid or specific siRNA. Lipofectamine 2000 was utilized for transfection.

### Statistical analysis

Statistical analyses were carried out in IBM SPSS 22.0. Student’s t-test (two-tailed, unpaired) served for two-group comparisons, and one-way ANOVA for three or more groups. Significance was set at *p* < 0.05. Figures were produced using GraphPad Prism 9.5.1.

## RESULTS

### Overview of the sequencing data

A total of 25 individual pigs covering three breeds were enrolled in this study, including 8 Duroc (DU1–DU8), 8 Jiangquan Black (JQ1–JQ8), and 9 Laiwu (LW1–LW9) pigs. Post-quality control sequencing metrics showed reliable data quality across all samples. For Duroc samples, the GC content fluctuated between 68.93% and 70.21%, with the Q30 value exceeding 92.16%. Approximately 80.87%–82.93% of clean reads were successfully aligned to the genome, among which uniquely mapped reads accounted for 45.86%–50.43% and multi-mapped reads occupied 25.88%–29.21% (Supplementary Table S3). For Jiangquan Black pig samples, GC content ranged from 49.90% to 53.77% and Q30 values were over 90.41%. These samples exhibited a higher genome mapping rate of 96.12%–97.60%, with 87.69%–89.12% unique mapped reads and merely 1.11%–3.22% multi-mapped reads (Supplementary Table S4). For Laiwu pig samples, the GC content and Q30 value were 52.76%–53.64% and above 90.03%, respectively. The overall mapping ratio was 91.46%–97.04%, consisting of 79.74%– 88.11% unique mapped reads and 4.16%–7.43% multi-mapped reads (Supplementary Table S5). Collectively, these quality indicators verified the high quality and credibility of the transcriptome sequencing data, which was qualified for subsequent bioinformatic analyses.

### Differential expression analysis

In the comparison between DU and JQ, there were a total of 5416 DE genes (2711 up- and 2706 down-regulated) (Figure 1A). In the comparison between DU and LW, there were a total of 4846 DE genes (3517 up- and 1329 down-regulated) (Figure 1B). In the comparison between JQ and LW, there were a total of 2777 DE genes (2499 up- and 278 down-regulated) (Figure 1C). Heatmap visualization demonstrated obvious inter-group expression differences of DEGs (Figure 1D, E, F). A total of 214 genes are shared among the three groups. (Figure 1G). Six DEGs were randomly picked for qRT-PCR to corroborate the RNA-seq findings. The high consistency between the two approaches across all groups (Supplementary Figure S1) validated the sequencing data quality. Among the DEGs, *CCND2* was selected for further functional validation based on the following criteria: (1) it was consistently upregulated in Duroc pigs in both DU vs. JQ and DU vs. LW comparisons; (2) it was significantly enriched in pathways known to regulate skeletal muscle development (PI3K-Akt, Wnt, and FoxO signaling); and (3) previous studies have reported its involvement in myogenesis. These lines of evidence suggested that *CCND2* may play a role in breed-specific muscle growth [31].

**Figure 1.**
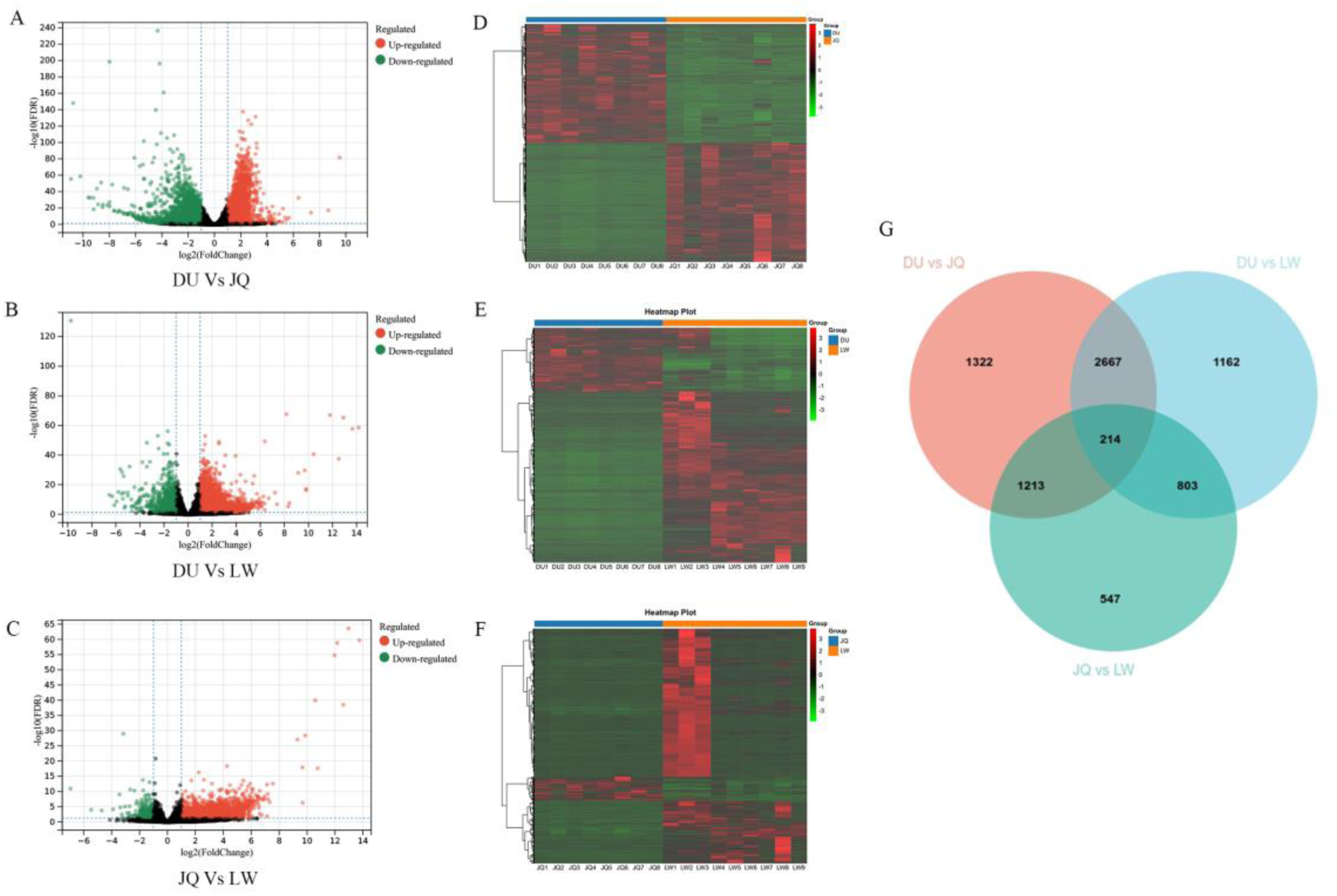
Statistics of DE mRNAs. (A, B, C) Volcano plot. Red notes represent upregulated genes, while green notes represent downregulated genes. (D, E, F) Heatmap of DE mRNAs. Rows: mRNAs; columns: samples. Red and green denote high and low expression levels, respectively (G) The Venn diagram of differential expression analysis.

### Weighted gene co-expression network analysis

A range of soft threshold was set from 1 to 30. When R^2^ = 0.9, the corresponding optimal soft threshold for this study was 16. The independence had a high degree and the average connectivity had lower level when the power value was equal to 16 (Figure 2A). Accordingly, a soft threshold power of 16 was adopted to construct the hierarchical clustering tree. Average linkage clustering partitioned 7938 genes (mean FPKM > 0.3) with akin expression patterns into different co-expression modules. Finally, eight modules (darkgreen, darkmagenta, darkorange, darkorange2, lightgreen, blue, florawhite, and grey) were identified and labeled with different colors (Figure 2B). The mRNA counts within each module were listed in Supplementary Table S6.

**Figure 2.**
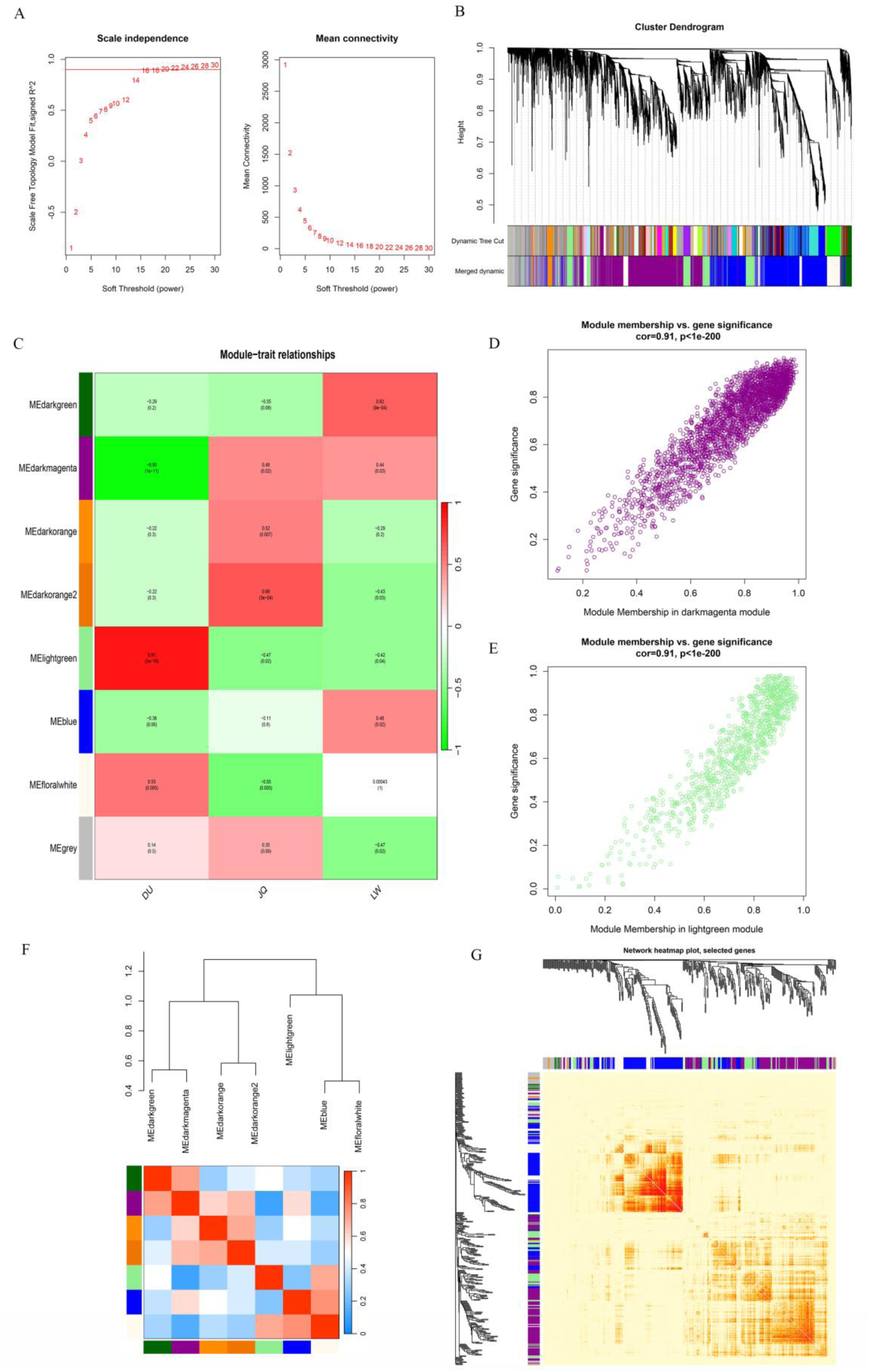
Statistics of WGCNA. (A)The selection of the optimal soft threshold.(B) The hierarchical clustering tree displays various modules and performs the merging of modules. (C) Heatmap of module-trait relationships. Colors denote correlation direction (red: positive; green: negative), and *p*-values are provided for each association. (D, E) GS Vs MM in the interested module. MM represents the correlation between genes and modules, while GS represents the correlation between genes and traits, and both are then correlated. (F) The correlation graph between various modules. (G) Visualization of gene network heatmap.

The correlations between co-expression modules and target traits were analyzed and visualized (Figure 2C). In this study, Duroc pigs were selected as the focal breed due to their rapid skeletal muscle growth rate, which was the core trait of interest. The grey module comprised unclassified genes that were not assigned to any specific co-expression module. The lightgreen module showed a strong positive correlation with Duroc pigs (*r* = 0.91, *p* = 5×10), but negative correlations with Jiangquan Black pigs (*r* = −0.47, *p* = 0.02) and Laiwu pigs (*r* = −0.42, *p* = 0.04). By contrast, the darkmagenta module exhibited an extremely significant negative correlation with Duroc pigs ( *r* = - 0.93, *p* = 1×10), together with positive correlations with Jiangquan Black pigs (*r* = 0.48, *p* = 0.02) and Laiwu pigs (*r* = 0.44, *p* = 0.03). In these two modules, Duroc pigs, which are the trait of interest in this study, exhibit a highly significant negative correlation with Jiangquan black pigs and Laiwu pigs. These two modules presented completely opposite correlation patterns between Duroc pigs and the two indigenous pig breeds, indicating that the genes within these modules are highly likely to contribute to skeletal muscle development.

In this study, correlation analysis between Gene significance (GS) and module membership (MM) was performed for the darkmagenta and lightgreen modules, respectively. The results demonstrated an extremely strong positive correlation between GS and MM in both modules (*cor* = 0.91, *p* < 1×10^-200^) (Figure 2D, E), confirming the high statistical significance and reliability of these two core modules. Furthermore, the correlation network among all co-expression modules was visualized (Figure 2F). Additionally, 400 genes were selected to construct a gene-gene correlation heatmap, and relatively strong correlations were clearly observed along the diagonal of the heatmap (Figure 2G).

### GO and KEGG enrichment analyses

To explore the functional relevance of breed-related DE genes and two muscle-associated modules, we conducted GO term and KEGG pathway enrichment analyses. This analysis yielded 75 BP, 78 CC, and 89 MF categories, alongside 98 KEGG pathways, with the top 20 terms presented in Figure 3A–D. In all significant GO terms and KEGG pathways, there were no directly associated BP with skeletal muscle growth and development. No significant GO terms directly related to muscle development were identified among the enriched categories. In contrast, multiple KEGG pathways with well-established roles in skeletal muscle growth and development reached significance, including PI3K-Akt (*p* = 7.44×10⁻⁴), Wnt (*p* = 0.002), Hippo (*p* = 0.005), insulin (*p* = 0.007), TGF-beta (*p* = 0.009), AMPK (*p* = 0.015), MAPK (*p* = 0.018), and FoxO (*p* = 0.025) signaling.

**Figure 3.**
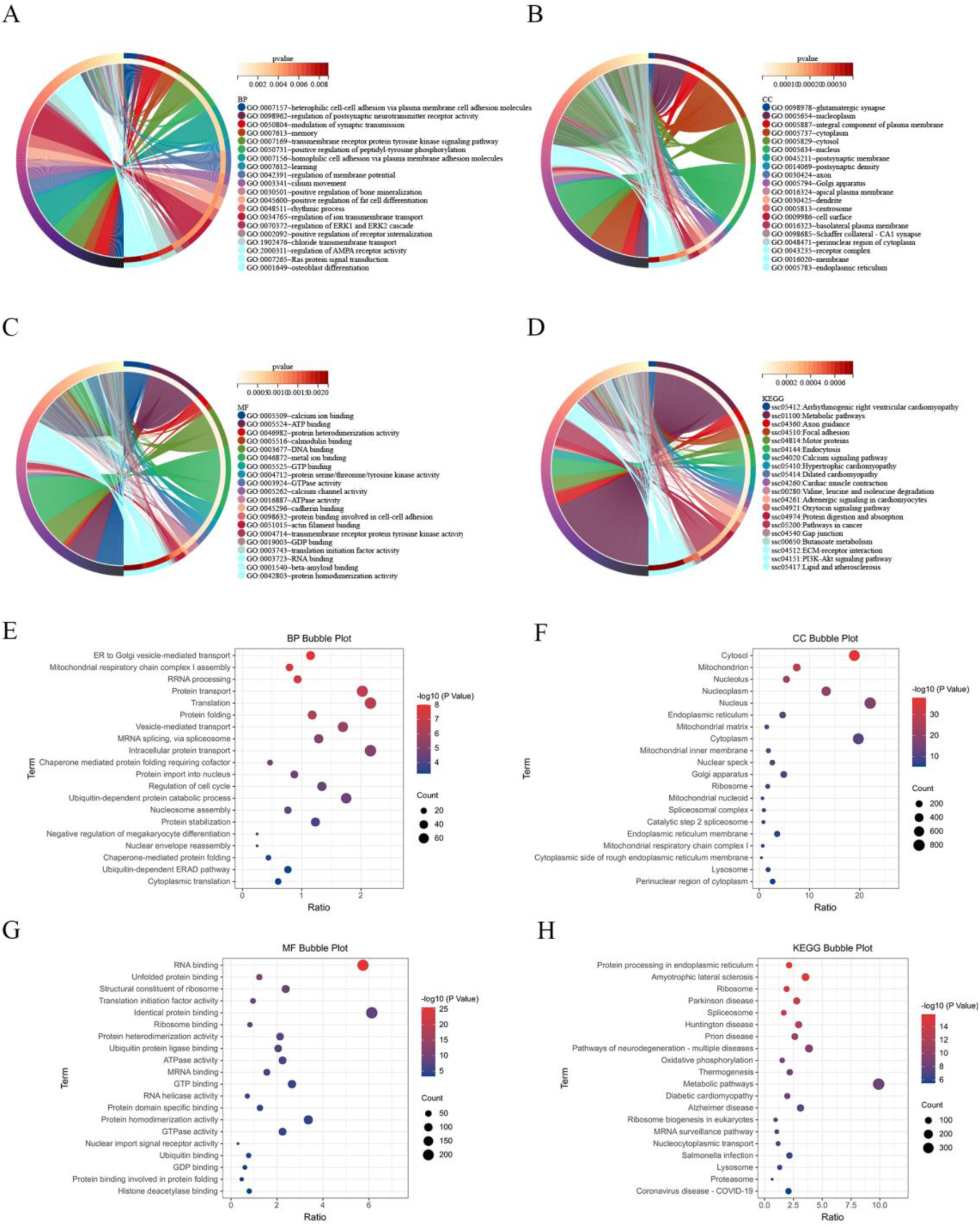
TOP 20 of GO enrichment analysis and KEGG pathways. Statistics of GO enrichment analysis and KEGG pathway for DE mRNA. (A) BP. (B) CC. (C) MF. (D) KEGG pathway. Statistical of GO enrichment analysis and KEGG pathway for two gene modules. (E) BP. (F) CC. (G) MF. (H) KEGG pathway.

In the present study, the genes of two modules were significantly enriched in 166 BP, 137 CC, 100 MF categories and 66 KEGG pathways. The top 20 results of the GO enrichment analysis and KEGG pathway analysis for the genes of two modules were shown (Figure 3E, F, G, H). Notably, significant muscle-relevant terms included skeletal muscle fiber development (*p* = 0.022), myoblast fusion (*p* = 0.026), insulin signaling (*p* = 8.48×10⁻⁴), and glucagon signaling (*p* = 0.020).

### The detection of C2C12 cell differentiation induction effect

To examine *CCND2* expression during C2C12 myogenesis, cells were differentiated and sampled every two days (D0–D8) for RNA extraction. qRT-PCR was performed to measure transcript levels of *CCND2* and the myogenic markers *MYOG*, *MEF2C*, and *MYF5*, with morphological changes recorded in Figure 4A and expression profiles in Figure 4B. Over the induction period, the cells progressively fused into multinucleated myotubes.

**Figure 4.**
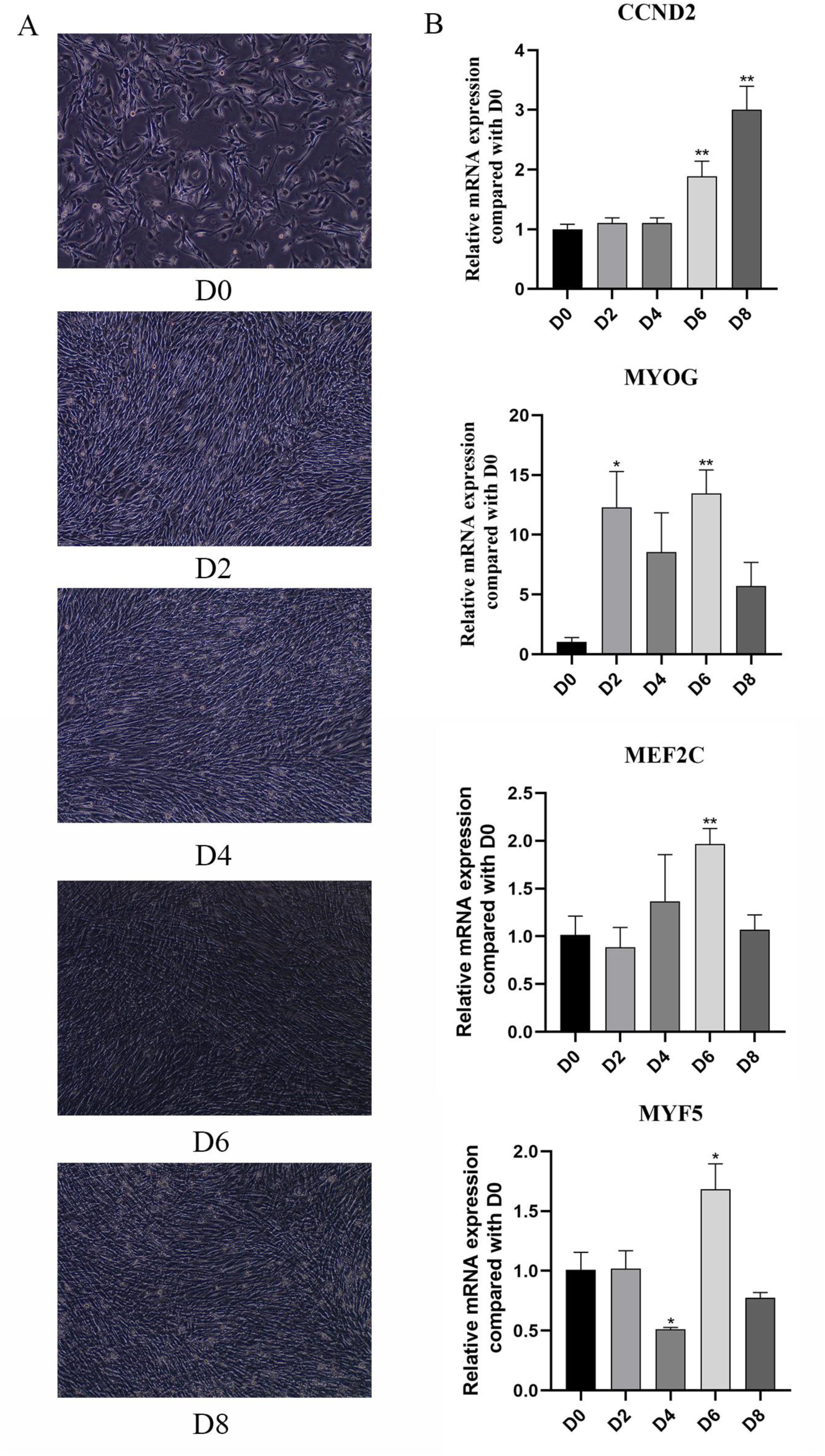
Detection of Induction differentiation effect of C2C12 cells. (A) Changes of cells during cell differentiation. (B) Expression levels of *CCND2*, *MYOG*, *MEF2C*, and *MYF5* during cell differentiation (The magnification factor of A is 4 times, * *p* < 0.05, \*\**p* < 0.01).

The expression level of *CCND2* showed no significant difference at D2 and D4 compared with D0, but was significantly upregulated from D6 onwards (*p*< 0.01) and reached the peak at D8 (*p* < 0.01). The myogenic differentiation markers *MYOG*, *MEF2C* and *MYF5* displayed an overall upward trend, with the maximum expression levels observed at D6. Therefore, D6 was selected as the time point for subsequent transfection and differentiation induction assays.

### Screening of interference fragments and detection of overexpression efficiency

Therefore, for the subsequent D6 transfection experiment, 3 μg of the overexpression vector pcDNA3.1(+)-CCND2 was chosen.

Four pairs of siRNAs targeting *CCND2* were designed and synthesized by Suzhou Genepharma Co., Ltd. C2C12 cells were transfected with 5 μl of negative control siRNA (si-NC) and four candidate *CCND2*-targeting siRNA fragments, respectively. After 24 h of incubation, total cellular RNA was extracted, and the mRNA expression level of *CCND2* was detected using qRT-PCR. The results revealed that only the si-385 fragment significantly downregulated *CCND2* expression (*p* < 0.05), indicating satisfactory interference efficiency, whereas the other three siRNA fragments exerted no significant silencing effect (Supplementary Figure S2). Accordingly, 5 μl of si-385 was selected for subsequent D6 transfection assays.

Following the manufacturer’s protocols, C2C12 cells were transfected with gradient doses (3 μg, 4 μg, 5 μg) of the empty pcDNA3.1(+) vector and the recombinant pcDNA3.1(+)-*CCND2* overexpression vector, respectively. Twenty-four hours post-transfection, total RNA was isolated from cells, and *CCND2* expression levels were quantified by qRT-PCR. The results demonstrated that all three plasmid doses significantly elevated *CCND2* expression (*p* < 0.001), with the most pronounced upregulation observed at a transfection dose of 3 μg (Supplementary Figure S2). Thus, 3 μg of the pcDNA3.1(+)-*CCND2* overexpression vector was chosen for follow-up D6 transfection experiments.

### The interference of *CCND2* inhibits myogenic differentiation of C2C12 cells

At D6 post-transfection, qRT-PCR analysis revealed that si-CCND2 treatment significantly reduced *CCND2*, *MYOG* and *MYF5* mRNA levels compared with si-NC controls (*p* < 0.05), whereas *MEF2C* transcripts showed no notable changes (Figure 5). The qRT-PCR results indicate that *CCND2* knockdown exerts an inhibitory effect on myogenic differentiation of C2C12 cells.

**Figure 5.**
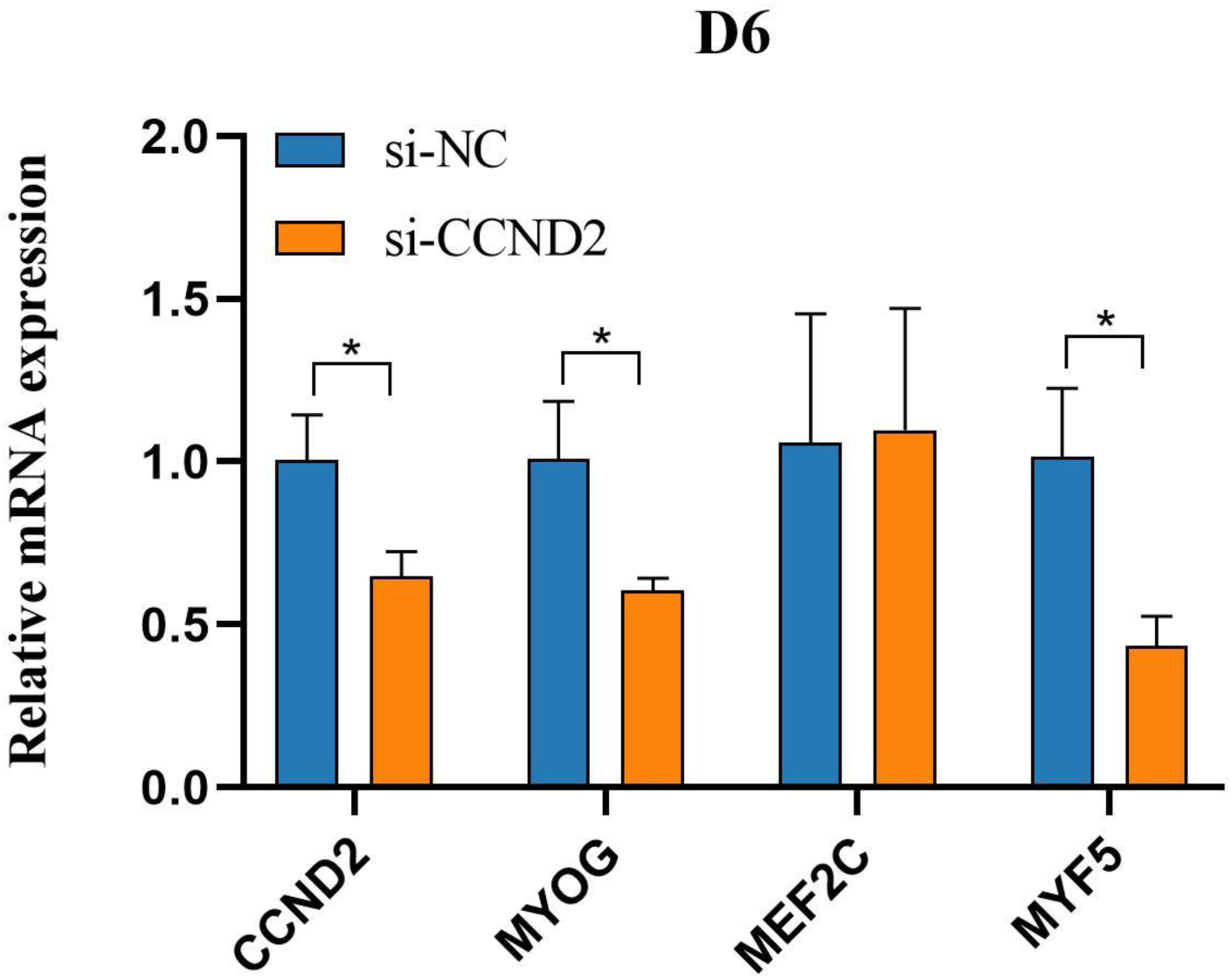
Knockdown of *CCND2* suppresses C2C12 differentiation. qRT-PCR for *CCND2*, *MYOG*, *MEF2C*, and *MYF5* at D6 (\**p* < 0.05).

### The overexpression of CCND2 promotes myogenic differentiation of C2C12 cells

At D6 post-transfection, qRT-PCR confirmed successful *CCND2* overexpression in pcDNA3.1(+)-CCND2-transfected cells compared with empty-vector controls. This upregulation was accompanied by significant increases in *MYOG* and *MEF2C* mRNA levels, whereas *MYF5* transcripts remained unchanged (Figure 6). The qRT-PCR results indicate that *CCND2* overexpression exerts a promoting effect on myogenic differentiation of C2C12 cells.

**Figure 6.**
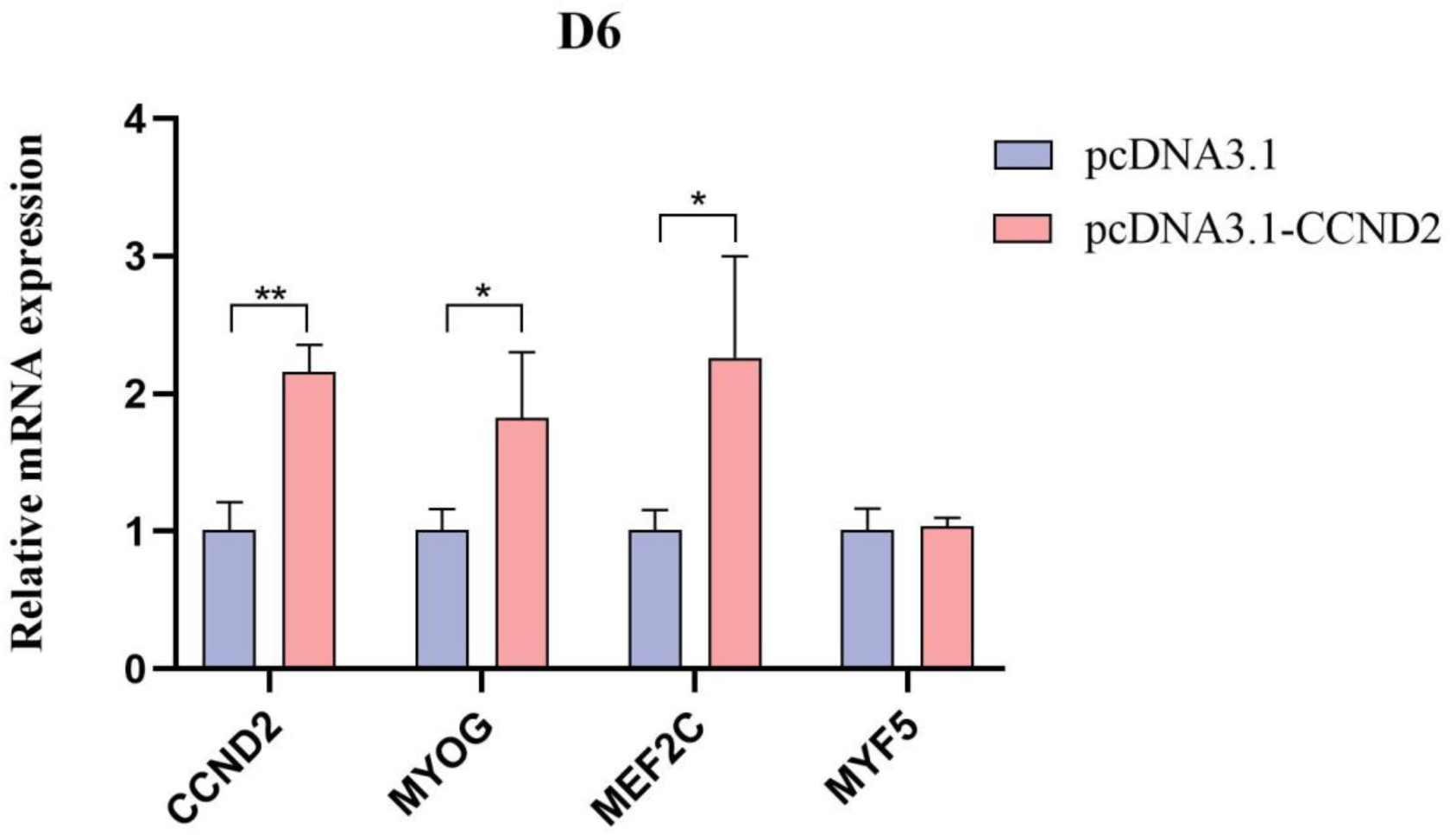
Overexpression of *CCND2* facilitates C2C12 differentiation. qRT-PCR for *CCND2*, *MYOG*, *MEF2C*, and *MYF5* at D6 (\**p* < 0.05, \**p* < 0.001).

## DISCUSSION

Porcine skeletal muscle is a highly heterogeneous tissue. As the most economically important and physiologically metabolically active skeletal muscle in pigs, the longissimus dorsi muscle has long been a key focus in molecular breeding research and genetic improvement programs [32]. Skeletal myogenesis encompasses a series of progressive events: commitment of muscle stem cells to mononucleated myoblasts, their alignment and fusion into multinucleated myotubes, and terminal maturation into functional myofibers [33–35]. This study seeks to provide insights for enhancing pig growth rate, improving meat production efficiency, and elevating pork quality.

To further elucidate the molecular mechanisms driving differential growth rates among distinct pig breeds, RNA-seq was performed on 25 samples in this study. The base quality score (Q) is calculated as Q = −10*log10(*p*) [36], where *p* represents the probability of an incorrect base call. In this study, the Q30% for all samples exceeded 89%, indicating that the sequencing data met the quality requirements and were suitable for further analysis. Each sample was aligned using Hisat2, and the overall alignment rate for each sample was above 80%, further supporting their suitability for subsequent analysis. For differential expression analysis, DEGs were screened using cutoffs of FDR < 0.05 and |log₂(FC)| ≥ 1. Pairwise comparisons were performed among three pig breeds: DU vs. JQ, DU vs. LW, and JQ vs. LW. The results showed that a greater number of DE genes were identified between Duroc—a well-known fast-growing breed—and the two local breeds (JQ and LW), whereas fewer DE genes were detected between JQ and LW.

IGF2 is known to positively regulate skeletal muscle differentiation [37]. Consistently, the IGF-2-encoding gene ENSSSCG00000035293 was enriched in our study in several muscle-related pathways, such as PI3K-Akt, insulin, and MAPK signaling. IGF-2 shows differential expression in DU vs. JQ and DU vs. LW, with high expression observed in Duroc pigs. Among these, Duroc pigs were identified as the fastest-growing breed in this study. Research has explored the role of FGF6 in skeletal muscle regeneration. FGF6-deficient mice exhibit notable regeneration defects post-injury, including fibrosis in muscle tissue and degeneration of skeletal muscle fibers. FGF6 deficiency in mice leads to impaired muscle regeneration, characterized by fibrosis, myofiber degeneration, and diminished MYOD/MYOG expression, confirming its pro-regenerative role [38]. Correspondingly, our dataset showed that FGF6 (ENSSSCG00000032662) was enriched in muscle-related pathways (PI3K-Akt and MAPK) and was more abundantly expressed in Duroc pigs relative to the other two breeds.

A study constructed co-expression networks and gene modules of abdominal aortic aneurysm (AAA) and control group expression data using WGCNA. Enrichment analysis of three modules identified 10 core genes that may play crucial roles in the pathogenesis of AAA, potentially offering diagnostic value [39]. Tian et al. applied WGCNA in breast cancer to construct gene co-expression networks, identifying significant modules and biomarkers. The study discovered four key genes associated with breast cancer prognosis and validated their potential roles in clinical aspects and targeted therapy of breast cancer [40]. Quan et al. applied WGCNA to pinpoint key immune-related genes in ovarian cancer. The high expression of these key genes correlates significantly with ovarian cancer prognosis and immune cell infiltration levels. Additionally, the study analyzed the functions of these key genes, revealing their involvement in T cell activation and differentiation, immune receptor activity [41]. WGCNA is suitable for studies involving more than 5 groups or 15 samples.

With a total of 25 samples in this study, WGCNA was deemed appropriate for the analysis. Duroc pigs, renowned globally for their high lean meat content and rapid growth, exhibit faster growth rates compared to Jiangquan and Laiwu pigs. Therefore,

Duroc pigs were selected as the focal breed for this WGCNA study. Analysis Module-trait relationships revealed that the lightgreen module displayed a strong positive correlation with Duroc pigs (cor = 0.91, *p* = 5×10^-5^), alongside negative correlation with Jiangquan and Laiwu pigs. Conversely, the darkmagenta module exhibits a strong negative correlation with Duroc pigs (cor = −0.93, *p* = 1×10^-11^), and positive correlation with Jiangquan and Laiwu pigs. These two modules, identified as key modules of interest in this study, showed strong and highly significant correlations with Duroc pigs, with an opposing trend observed in Jiangquan and Laiwu pigs. This suggests that within these two gene modules, gene expression levels in Duroc pigs are negatively correlated with those in Jiangquan and Laiwu pigs, indicating a potential role in skeletal muscle formation.

Shi et al. demonstrated that knocking out the MYMK gene in zebrafish embryos results in defective myofiber fusion in fast muscle, reduced muscle growth, and increased infiltration of fat cells [42]. In the present study, the gene ENSSSCG00000022891 (MYMK) was identified as a member of the darkmagenta module and was significantly enriched in the BP of myoblast fusion involved in muscle regeneration. The abundance of ENSSSCG00000022891 (MYMK) in Duroc pigs (mean FPKM = 0.59) is higher compared to Jiangquan pigs (mean FPKM = 0.28) and Laiwu pigs (mean FPKM = 0.36). This finding is consistent with the established knowledge that Duroc pigs exhibit the fastest growth rate among these three breeds. Wang et al. demonstrated that silencing the IDE gene significantly enhanced the survival and quantity of skeletal muscle stem cells. This suggests that the IDE gene negatively regulates skeletal muscle development, potentially inhibiting muscle growth [43]. In our research, ENSSSCG00000010470 (IDE) was identified as a member of the lightgreen module. The expression level of ENSSSCG00000010470 (IDE) was lower in Duroc pigs compared to Jiangquan pigs and Laiwu pigs. This corresponds to scientific knowledge that Duroc pigs exhibit the fastest growth rate among these three pig breeds.

Myogenic differentiation is essential for muscle fiber formation [44]. In our analysis, *CCND2* transcript abundance was significantly higher in Duroc pigs than in JQ and LW animals, and the gene was overrepresented in FoxO, PI3K-Akt, and Wnt cascades—pathways with established roles in muscle development. Given the limited evidence regarding *CCND2*’s involvement in myogenic differentiation, we examined its functional contribution through loss- and gain-of-function approaches in C2C12 cells. qRT-PCR analyses demonstrated that *CCND2* overexpression at D6 significantly elevated its own transcript alongside myogenic markers (*p* < 0.05), whereas *CCND2* silencing produced the opposite effect (*p* < 0.05). These results establish *CCND2* as a positive regulator of C2C12 myogenic differentiation. This conclusion is further supported by Zhou et al [31], who reported that *MyoD1* may modulate *CCND2* expression through the PI3K-Akt axis, thereby influencing muscle formation in Guangling cattle; they also found that *MyoD1* knockdown reduces *CCND2* transcript levels, reinforcing its role as a downstream effector in myogenic differentiation.

RNA-seq was performed to screen for DEGs among pig breeds exhibiting distinct growth rates, and the role of *CCND2* in C2C12 cell differentiation was examined. Nevertheless, this study has several limitations. It was limited to identifying DEGs among pig breeds with different growth rates at the transcriptomic level and specifically focused on investigating the role of *CCND2* in muscle cell differentiation using the C2C12 cell. Advanced sequencing technologies and improved experimental approaches are needed to further elucidate the role of *CCND2* in skeletal muscle development.

## CONCLUSION

In conclusion, this study conducted transcriptome analysis on Jiangquan Black pigs, Duroc pigs, and Laiwu pigs, identifying DEGs among these breeds. WGCNA identified two distinct gene modules potentially associated with growth rate. The DEGs and two module genes showed significant enrichment in BP terms and KEGG pathways linked to muscle development. Functional experiments involving *CCND2* knockdown and overexpression in C2C12 cells demonstrated that *CCND2* promotes myogenic differentiation. Collectively, these results identify the genetic basis for pork quality and skeletal muscle development, offering key references for pig molecular breeding to boost growth and meat quality.

## CONFLICT OF INTEREST

No potential conflict of interest relevant to this article was reported.

## AUTHORS’ CONTRIBUTION

Conceptualization: Chen W, Zeng YQ

Data curation: Li JB, Zhang JM

Formal analysis: Li JB, Song Q

Methodology: Li JB, Li SY

Software: Li JB, Li SY

Validation: Li JB, Zhang JM

Investigation: Li JB, Song Q

Writing - original draft: Li JB

Writing - review & editing: Li JB, Chen W

## FUNDING

This research was funded by the financial support provided by the Shandong Provincial Natural Science Foundation of China (ZR2022MC065).

## ACKNOWLEDGMENTS

We are grateful to Shiyin Li, Yingbin Du, Jianmin Zhang, and Qi Song for their assistance in sample collection and laboratory analyses.

## SUPPLEMENTARY MATERIAL

Supplementary file is available from:

## DATA AVAILABILITY

The transcriptomic data are available in the Sequence Read Archive of the NCBI under the accession number PRJNA1482246.

## DECLARATION OF GENERATIVE AI

No AI tools were used in this article.

## Notes

### Competing Interest Statement

The authors have declared no competing interest.

