## Supplementary Material for "Transcriptome analysis reveals *CCND2* promotes myogenic differentiation of pig skeletal muscle"

Supplementary Table S1. The qRT-PCR primer sequences

| 基因<br>Gene | 序列<br>Sequence (5’-3’) |
|---|---|
| Sus-GAPDH-F | AAGTTCCACGGCACAGTCAAG |
| Sus-GAPDH-R | CACCAGCATCACCCCATTT |
| Sus- ROMO1-F | GCTTTGACCGCGTGAAGATG |
| Sus-ROMO1-R | CGCATTCCGATCCTGAGACA |
| Sus-KLF13-F | GGATCCTGGCGGACCTTAAC |
| Sus-KLF13-R | GGTGCGAGGATTTCCCGTAA |
| Sus-NT5DC2-F | CCAACTCTTCGACGTGGTCA |
| Sus-NT5DC2-R | TTTTCTGAAAGGCTTGCGCC |
| Sus-SLC25A47-F | GGTCTCGTCCGTGTCTTTCG |
| Sus-SLC25A47-R | TCAGCACCACTTCATAGGCG |
| Sus-TMEM140-F | CTACACGTGGAAGTGGCTCA |
| Sus-TMEM140-R | GAGGATGAGTAAGGCCTGGG |
| Sus-CYP2A19-F | ACTGAACCTCTTCTTCGCGG |
| Sus-CYP2A19-R | AACTGCCCGTTCTCATCCAG |
| Mus-β-actin-F | GTGACGTTGACATCCGTAAAGA |
| Mus-β-actin-R | GCCGAACTCATCGTACTCC |
| Mus-CCND2-F | AGTCCCGACTCCTAAGACCC |
| Mus-CCND2-R | GCGTTATGCTGCTCTTGACG |
| Mus-MYOG-F | GAGACATCCCCCTATTTCTACCA |
| Mus-MYOG-R | GCTCAGTCCGCTCATAGCC |
| Mus-MEF2C-F | AAATCTCTCCCTGCCTTCTA |
| Mus-MEF2C-R | GGTGTGTTGTGGGTATCTCG |
| Mus-MYF5-F | TCAGACGATGAGGAGCACG |
| Mus-MYF5-R | GAGTCTCTGGTTAGGGTTGGTG |

Supplementary Table S2. Sequence information of siRNA interference

| 基因<br>Gene | 上游引物<br>Forward(5'-3') | 下游引物<br>Forward(5'-3') |
|---|---|---|
| CCND2-Mus-385 | CCGCAGUGUUCCUAUUUCATT | UGAAAUAGGAACACUGCGGTT |
| CCND2-Mus-599 | CCAAGCUGAAAGAGACCAUTT | AUGGUCUCUUUCAGCUUGGTT |
| CCND2-Mus-841 | GACUUCAAGUUUGCCAUGUTT | ACAUGGCAAACUUGAAGUCTT |
| CCND2-Mus-918 | GGAUGAUGAAGUGAACACATT | UGUGUUCACUUCAUCAUCCTT |
| si-NC | UUCUCCGAACGUGUCACGUTT | ACGUGACACGUUCGGAGAATT |

Supplementary Table S3. Overview of the Duroc pigs sequencing data

| | DU1 | DU2 | DU3 | DU4 | DU5 | DU6 | DU7 | DU8 |
|---|---|---|---|---|---|---|---|---|
| Total Reads Counts(#) | 120013800 | 130792768 | 112828462 | 106306746 | 111629760 | 125421664 | 109355724 | 113830698 |
| Total Bases Counts(bp) | 17850996854 | 19445712704 | 16773174978 | 15797988718 | 16578208270 | 18614868072 | 16244588426 | 16900149010 |
| Average read length(bp) | 148 | 148 | 148 | 148 | 148 | 148 | 148 | 148 |
| Q30 Bases Count(bp) | 16452391824 | 18016037750 | 15560314649 | 14644497783 | 15296868014 | 17190848314 | 14998229426 | 15633741519 |
| Q30 Bases Ratio(%) | 92.16 | 92.64 | 92.76 | 92.69 | 92.27 | 92.35 | 92.32 | 92.50 |
| GC Bases Ratio(%) | 69.67 | 69.44 | 69.29 | 69.36 | 68.93 | 70.21 | 70.21 | 69.60 |
| Uniquely mapped reads | 57426544 | 60866346 | 56896058 | 52232782 | 55593824 | 59296384 | 50147998 | 54126462 |
| % of Uniquely mapped reads | 47.85 | 46.54 | 50.43 | 49.13 | 49.80 | 47.28 | 45.86 | 47.55 |
| Mutiple mapped reads | 34117700 | 38210888 | 29203188 | 29416932 | 29856764 | 34654160 | 31536668 | 31998448 |
| % of Mutiple mapped | 28.43 | 29.21 | 25.88 | 27.67 | 26.75 | 27.63 | 28.84 | 28.11 |
| unmapped reads | 28128066 | 31201412 | 26271938 | 24293926 | 25922684 | 31093322 | 27377270 | 27394956 |
| % of unmapped reads | 23.44 | 23.86 | 23.28 | 22.85 | 23.22 | 24.79 | 25.04 | 24.07 |
| Overall alignment rate (%) | 82.78 | 82.93 | 82.71 | 82.46 | 81.66 | 81.11 | 80.87 | 82.01 |

Supplementary Table S4. Overview of the Jiangquan black pigs sequencing data

| | JQ1 | JQ2 | JQ3 | JQ4 | JQ5 | JQ6 | JQ7 | JQ8 |
|---|---|---|---|---|---|---|---|---|
| Total Reads Counts(#) | 45748468 | 41076582 | 53171760 | 45408790 | 45673282 | 51727794 | 47425616 | 46199494 |
| Total Bases Counts(bp) | 6843938453 | 6148760056 | 7957469642 | 6795659735 | 6835558810 | 7738313408 | 7099404109 | 6915183984 |
| Average read length(bp) | 149 | 149 | 149 | 149 | 149 | 149 | 149 | 149 |
| Q30 Bases Count(bp) | 6217910182 | 5559585268 | 7562092088 | 6458139917 | 6525018423 | 7168929375 | 6776661111 | 6581383933 |
| Q30 Bases Ratio(%) | 90.85 | 90.41 | 95.03 | 95.03 | 95.45 | 92.64 | 95.45 | 95.17 |
| GC Bases Ratio(%) | 53.43 | 52.73 | 53.63 | 53.76 | 53.77 | 49.90 | 53.50 | 52.66 |
| Uniquely mapped reads | 40168986 | 35831166 | 46624570 | 40157336 | 40190370 | 45573376 | 42265682 | 41038686 |
| % of Uniquely mapped reads | 87.80 | 87.23 | 87.69 | 88.44 | 88.00 | 88.10 | 89.12 | 88.83 |
| Mutiple mapped reads | 156232 | 112416 | 193896 | 158332 | 180040 | 76768 | 148558 | 150094 |
| % of Mutiple mapped | 2.54 | 1.96 | 2.93 | 3.22 | 3.22 | 1.11 | 3.13 | 2.96 |
| unmapped reads | 3073688 | 2862606 | 3310380 | 2457024 | 2791432 | 3450378 | 2372632 | 2534296 |
| % of unmapped reads | 6.72 | 6.97 | 6.23 | 5.41 | 6.11 | 6.67 | 5.00 | 5.49 |
| Overall alignment rate (%) | 96.52 | 96.31 | 97.02 | 97.30 | 97.04 | 96.12 | 97.60 | 97.38 |

Supplementary Table S5. Overview of the Laiwu pigs sequencing data

| | LW1 | LW2 | LW3 | LW4 | LW5 | LW6 | LW7 | LW8 | LW9 |
|---|---|---|---|---|---|---|---|---|---|
| Total Reads Counts(#) | 116108644 | 113823740 | 116322536 | 34277080 | 33792938 | 32751676 | 42073208 | 35623318 | 35665076 |
| Total Bases Counts(bp) | 11577775150 | 11351536168 | 11597808122 | 4308307269 | 4246370865 | 4114712963 | 5287083501 | 4476559914 | 4482251747 |
| Average read length(bp) | 99 | 99 | 99 | 125 | 125 | 125 | 125 | 125 | 125 |
| Q30 Bases Count(bp) | 10536997805 | 10298397891 | 10563213457 | 3883824774 | 3827480374 | 3692260031 | 4768140152 | 4005625290 | 4035764473 |
| Q30 Bases Ratio(%) | 91.01 | 90.72 | 91.07 | 90.14 | 90.13 | 89.73 | 90.18 | 89.47 | 90.03 |
| GC Bases Ratio(%) | 53.31 | 53.03 | 52.80 | 52.91 | 53.19 | 53.64 | 53.09 | 52.76 | 53.32 |
| Uniquely mapped reads | 102301054 | 98844062 | 101719280 | 28006772 | 27781062 | 26117184 | 33997350 | 28689046 | 28899316 |
| % of Uniquely mapped reads | 88.11 | 86.84 | 87.45 | 81.71 | 82.21 | 79.74 | 80.81 | 80.53 | 81.03 |
| Mutiple mapped reads | 6974346 | 8451992 | 7936088 | 1666860 | 1708510 | 1673832 | 1957354 | 1480550 | 1680676 |
| % of Mutiple mapped | 6.01 | 7.43 | 6.82 | 4.86 | 5.06 | 5.11 | 4.65 | 4.16 | 4.71 |
| unmapped reads | 6373258 | 6043700 | 6114430 | 4365454 | 4085540 | 4690630 | 5761336 | 5178038 | 4782450 |
| % of unmapped reads | 5.49 | 5.31 | 5.26 | 12.74 | 12.09 | 14.32 | 13.69 | 14.54 | 13.41 |
| Overall alignment rate (%) | 96.88 | 96.95 | 97.04 | 92.43 | 92.84 | 91.46 | 92.01 | 91.53 | 92.18 |

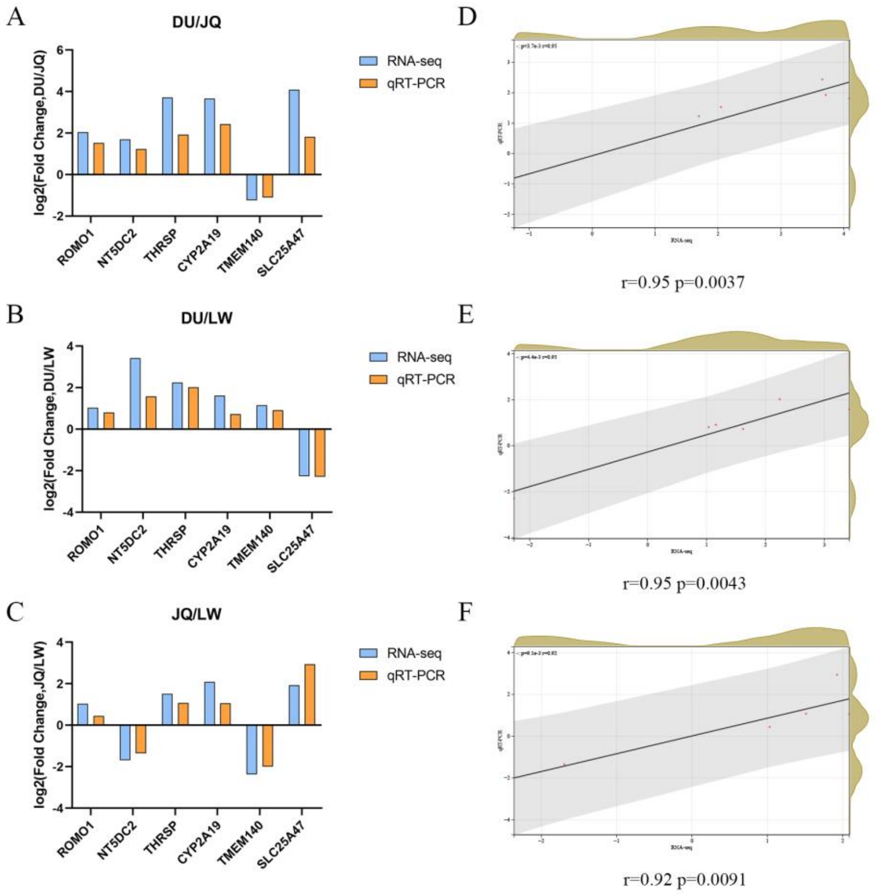


Supplementary Fig. S1 Validation by qRT-PCR of 6 randomly selected DE genes from RNA-seq. (A, B, C) Validation of qRT-PCR (D, E, F) Correlation and significance of RNA-seq and qRT-PCR

Supplementary Table S6. The number of genes in the 8 modules.

| Module | number |
|---|---|
| blue | 2502 |
| darkgreen | 206 |
| darkmagenta | 2678 |
| darkorange | 126 |
| darkorange2 | 37 |
| florawhite | 755 |
| lightgreen | 980 |
| grey | 654 |

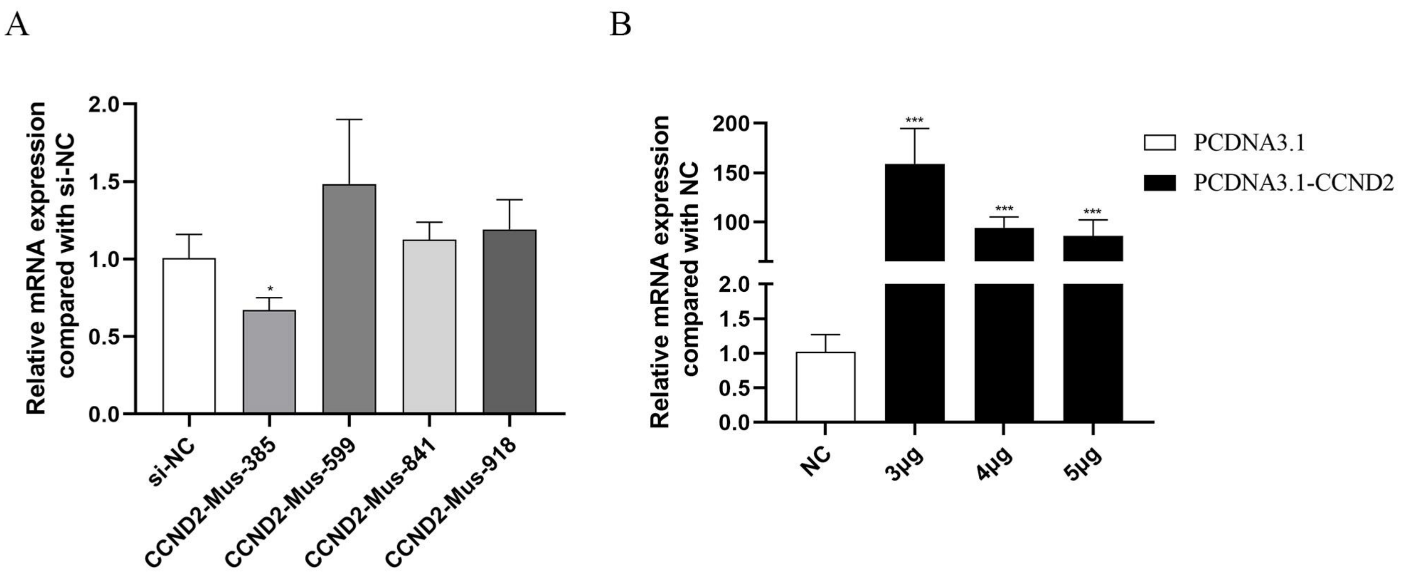


Supplementary Fig. S2 Statistics of transfection efficiency. (A) The relative expression level of CCND2 after transfection with the interfering fragment. (B) The relative expression level of CCND2 after transfection with the overexpression vector.
